# An alternative malonyl-CoA producing pathway in nature

**DOI:** 10.1101/2022.10.28.514148

**Authors:** Bo Liu, Yuwei Zhang, Qianqian Cui, Sheng Wu, Shuangyan Tang, Yihua Chen, Yanhe Ma, Weifeng Liu, Yong Tao

## Abstract

Malonyl-CoA is a key metabolic intermediate for biosynthesis of diverse cellular molecules and natural products. Carboxylation of acetyl-CoA is almost the unique pathway for malonly-CoA biosynthesis. Biotechnological production of numerous value-added malonyl-CoA-derived chemicals require high intracellular supply of malonyl-CoA. However, because of the central role of acetyl-CoA in primary metabolism, it is difficult to develop flexible strategies to balance malonyl-CoA supply with other cellular metabolism. Here we find that there is a natural alternative malonyl-CoA-producing pathway, in which the key reaction is catalyzed by an α-keto acid dehydrogenase complex BkdFGH from *Streptomyces avermitilis*. This dehydrogenase complex could efficiently catalyze biosynthesis of malonyl-CoA from oxaloacetate in addition to recognizing its native substrate branched-chain α-keto acid. Oxaloacetate dehydrogenase (OADH) was shown to play important physiological roles during the regulation of biosynthesis of native malonyl-CoA-derived polyketides in *Streptomyces*. Furthermore, the oxlaocetate dehydrogenation reaction is thermodynamically superior to acetyl-CoA carboxylation and enable efficient bioproduction of diverse malonyl-CoA-derived chemicals in engineering *Escherichia coli*. This novel malonyl-CoA source thus has great potential in the biotechnological field.

---

Malonyl-CoA is among the most important building blocks in cellular metabolism and biotechnological application^1, 2^. It is the essential precursors of diverse chemicals, including long-chain fatty acids, organic acids, polyketides, phenylpropanoids, and flavonoids (Fig. 1a), which have a wide range of applications in medicine, health care, and agriculture ^3-5^. Meanwhile, malonyl-CoA is also a key signaling molecule and participates in diverse range of physiological or pathological, and its intracellular level is thus tightly controlled. In the biotechnological field, improvement of malonyl-CoA is still a particularly challenging task. It is generally acknowledged that malonyl-CoA is produced almost exclusively by acetyl-CoA carboxylase ^1, 6, 7^, which catalyzes the formation of malonyl-CoA from acetyl-CoA and bicarbonate upon ATP consumption ^8^ (Δ_r_G^’m^ = +38.1 kJ/mol according to eQuilibrator). In *Escherichia coli*, intracellular malonyl-CoA is involved in cell growth and its abundance is controlled to a very low level ^9-11^, which is problematic for its heterologous biosynthesis.

**Fig. 1.**
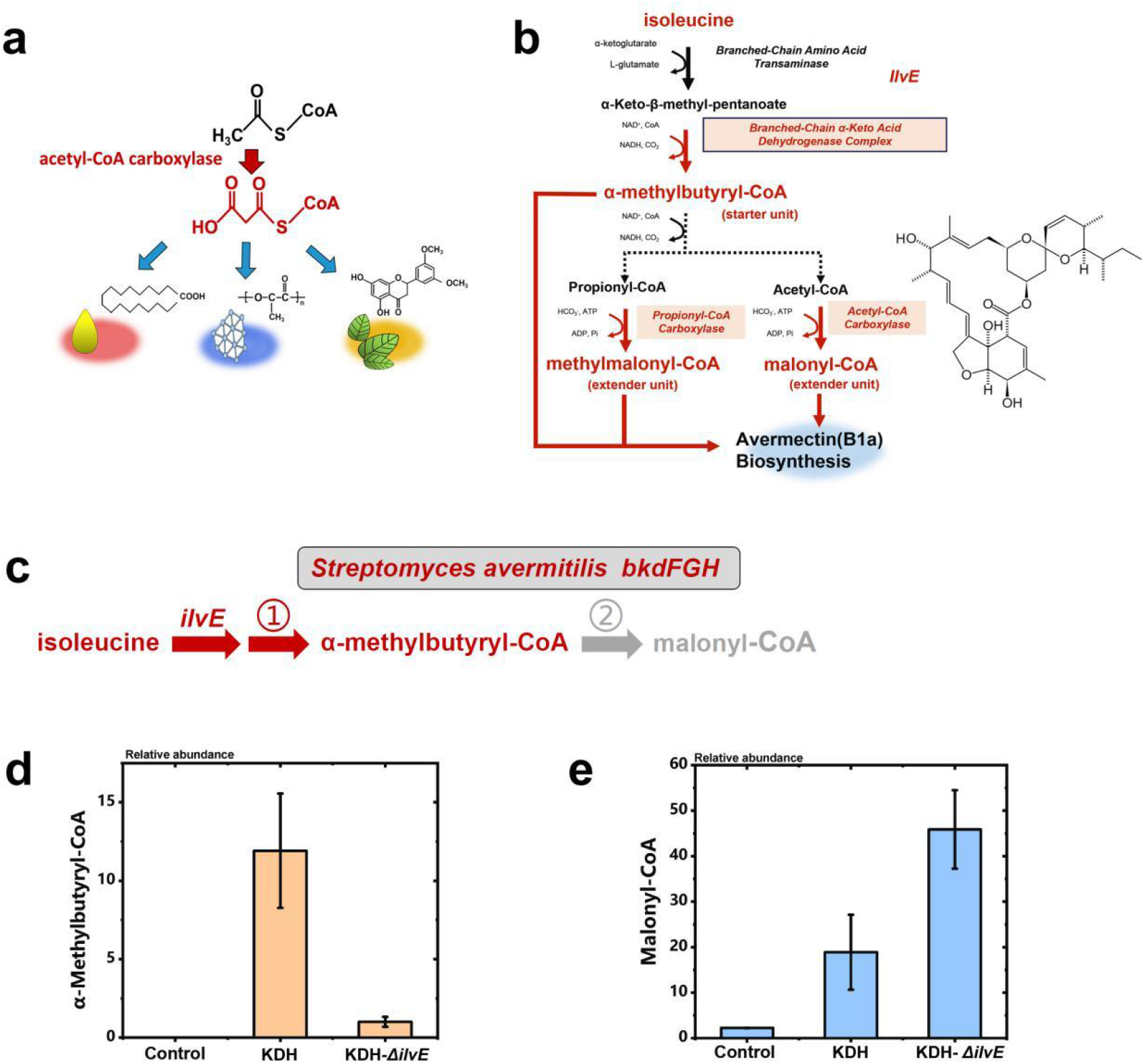
Serendipic increase of intracellular malonyl-CoA levels by α-keto acid dehydrogenase complex. **a**, Acetyl-CoA provides the major source of malonyl-CoA for biosynthesis of malonyl-CoA-derived chemicals. **b**, Metabolic engineering strategy to co-produce the three precursors of avermectin B1a aglycon, malonyl-CoA, α-methylbutyryl-CoA, and methylmalonyl-CoA from isoleucine. **c**, Expression of *Streptomyces avermitilis bkdFDH* lead to the increase of both α-methylbutyryl-CoA and malonyl-CoA levels. A possible metabolic route of malonyl-CoA and α-methylbutyryl-CoA biosynthesis is shown. **d-e**, Intracellular metabolite abundances of different strains. Relative abundance of α-methylbutyryl-CoA (**d**) and malonyl-CoA (**e**) for the Control (*Escherichia coli* BW25113 strain), KDH, and KDH-*ΔilvE* strains.

In recent decades, bioengineering strategies to improve intracellular malonyl-CoA levels are almost completely dealing with acetyl-CoA carboxylation pathway. These include mining novel acetyl-CoA carboxylases with higher activities ^12, 13^; rational metabolic network modification or systematic tuning carbon flux toward malonyl-CoA ^1, 6^; downregulating endogenous competitive malonyl-CoA consumption pathways ^2, 14, 15^. However, the complex regulation of acetyl-CoA metabolism and low efficiency of biotin-dependent carboxylase are the major impeding factors of the metabolic engineering for malonyl-CoA supply. Acetyl-CoA is a key “hub” molecule in central metabolic networks, important precursor of fatty acids as well as amino acids, and also indirectly involved in cellular energy metabolism through TCA cycle and oxidative phosphorylation.

Because the metabolic profiles are various in different engineering cells, it is difficult to develop a broadly effective strategy to balance malonyl-CoA biosynthsis and cellular metabolism. Furthermore, aceytl-CoA carboxylation is an energy-consuming reaction ^7^, and overexpression of acetyl-CoA carboxylase might affect the cell viability ^1^, which limit the modification effect of acetyl-CoA carboxylase in metabolic engineering.

We herein present an alternative malonyl-CoA-producing pathway in which the key reaction is catalyzed by the α-keto acid dehydrogenase complex BkdFGH from *Streptomyces avermitilis*. This complex is typically employed as the branched-chain α-keto acid dehydrogenase complex that supplies branched-chain acyl-CoA for avermectin biosynthesis during stationary phase ^16, 17^. However, when overexpressed in *E. coli*, we found that the intracellular malonyl-CoA level is dramatically increased by serendipity and independent of intracellular acetyl-CoA level. α-Keto acid dehydrogenase complex is an enzyme family that catalyze the oxidative decarboxylation of different α-keto acids, including branched-chain α-keto acids, pruvate, and α-ketogulatrate, and produce branched-chain-acyl-CoA, acetyl-CoA, and succinyl-CoA, respectively. We hypothesized malonyl-CoA produced by BkdFGH is generated from oxaloacetate, a common intermediate of tricarboxylic acid cycle (TCA cycle), through a similar catalytic mechanism. Our metabolic and biochemical studies confirmed this hypothesis. The BkdFGH complex has oxaloacetate dehydrogenase (OADH) activity (ΔrG’m = -99.0 kJ/mol according to eQuilibrator) and enable efficient malonyl-CoA using oxaloacetate as substrate both *in vivo* and *in vitro*. Nevertheless, comparative transcriptome reveals that the OADH in *Streptomyces* is required in both the cell-growth and polyketides accumulation periods. Thus, we proposed biosynthesis of malonyl-CoA by dehydrogenation from oxlaocetate is a natural biochemical reaction and an important alternative pathway for malonyl-CoA supply.

Taking the advantage of high reaction efficiency, dehydrogenation from oxaloacetate provides a promising alternative malonyl-CoA biosynthesis pathway for biotechnological purpose. Accordingly, this novel malonyl-CoA source was evaluated for the biosynthesis of diverse malonyl-CoA-derived products, including 3-hydroxypropionate, triacetate lactone, phloroglucinol, aloesone, flaviolin, raspberry ketone, resveratol, the titers of which were increased significantly. Thus, our findings present a novel malonyl-CoA source appropriate for the biosynthesis of value-added chemicals such as polyketides, flavonoids, and fatty acids in a cost-insensitive but efficiency-sensitive manner.

## Results

### Efficient malonyl-CoA production by α-keto acid dehydrogenase BkdFGH through a non-isoleucine pathway

Enhancing the activity of acetyl-CoA carboxylase is the major metabolic engineering strategy to improve malonyl-CoA supply. In our previous study, to develop potential chassis cells for the heterogeneous production of avermectin B1a, we designed a metabolic engineering strategy to co-produce the three acyl-CoA precursors of avermectin B1a aglycon, malonyl-CoA, α-methylbutyryl-CoA, and methylmalonyl-CoA. In this strategy, all three acyl-CoA precursors are coordinate produced from isoleucine pathway by enhancing branched-chain α-keto acid dehydrogenase complex, propionyl-CoA carboxylase, and acetyl-CoA carboxylase (Fig. 1b). However, when we expressed the branched-chain α-keto acid dehydrogenase complex (BCKDH) *bkdFGH* genes from *S. avermitilis* (MA-4680) to supply the starter unit α-methylbutyryl-CoA for avermectin B1a, accompanied by the increase of α-methylbutyryl-CoA level, malonyl-CoA level also increased approximately 8.6-fold even without enhancement of acetyl-CoA carboxylase (strain KDH, Fig. 1d-e) ^16, 17^.

We suspected if the extra malonyl-CoA is still generated through our purposed pathway by acetyl-CoA carboxylation (Fig. 1c), in which the precursor is provided by the dissociation of α-methylbutyryl-CoA. To test this hypothesis, we knocked out the branched-chain amino acid transaminase gene *ilvE* of strain KDH (obtaining strain KDH-Δ*ilvE*) to block the biosynthesis of α-keto-β-methylvaleric acid ^18^, a natural substrate of branched-chain α-keto acid dehydrogenase ^19^. As expected, the α-methylbutyryl-CoA level decreased dramatically (by 92%) (Fig. 1d). However, the malonyl-CoA level unexpectedly increased by 143%, a 21-fold increase over the control (Fig. 1e). This surprising result shows that the hypothesis above is invalid and reveals that the increase in malonyl-CoA level is unrelated to the acetyl-CoA carboxylation pathway.

### Oxaloacetate sustains efficient malonyl-CoA production by α-keto acid dehydrogenase

Given the results from the above experiments and the known catalytic mechanism of branched-chain α-keto acid dehydrogenase, we proposed a bold hypothesis that the malonyl-CoA is produced by directly dehydrogenation of BkdFGH, with the most likely substrate oxaloacetate (Fig. 2a). Therefore, we introduced BkdFGH into an oxaloacetate supply-enhancing strain. Phosphoenolpyruvate carboxylase (*ppc*) from *Corynebacterium glutamicum* was overexpressed in *E. coli* to supply more oxaloacetate (Fig. 2b). Strain KDH-PPC was obtained by integrating the *ppc* gene into the strain KDH genome (*ompT* loci) with a strong constitutive promoter P_tac_. As expected, the intracellular malonyl-CoA level was further increased 1.9-fold (strain KDH-PPC, Fig. 2c). We also compared the intracellular malonyl-CoA, acetyl-CoA, and oxaloacetate levels of strain KDH treated with glucose or oxaloacetate (Fig. 2d-f). To prevent possible interference from intracellular metabolism, cells were starved prior to treatment. The two experimental groups were named OAA (oxaloacetate treated) and Glc (glucose treated). After starvation for 2 h and oxaloacetate or glucose treatment for 1 h, the acetyl-CoA level of the OAA group was just 1.6% that of the Glc group, while the malonyl-CoA level was approximately 2.1-fold higher and the intracellular oxaloacetate level was 5.7-fold higher. The above data strongly indicate that oxaloacetate promotes the production of intracellular malonyl-CoA, and that the increased malonyl-CoA production is independent of acetyl-CoA level for strain KDH.

**Fig. 2.**
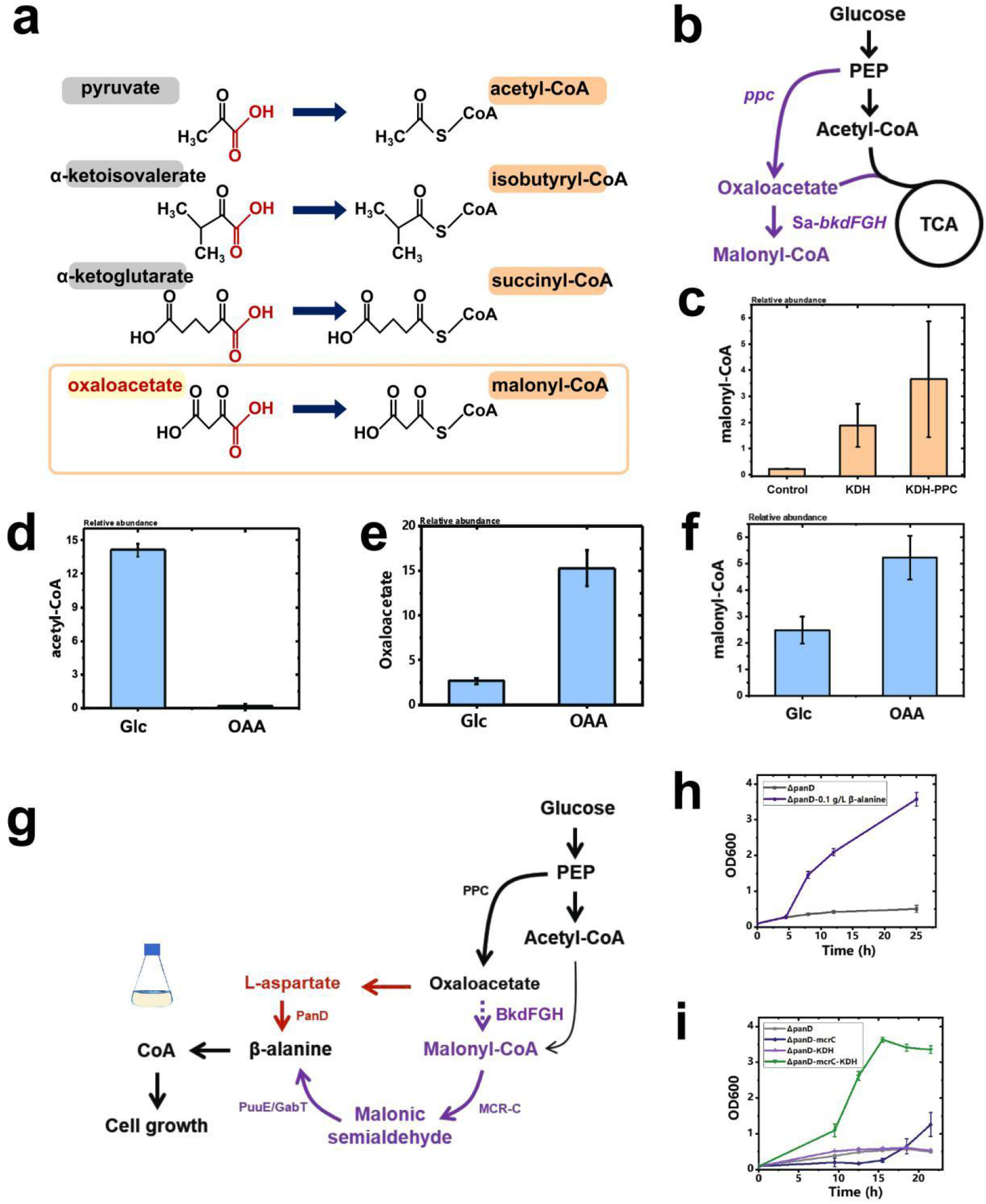
α-Keto acid dehydrogenase complex increases intracellular malonyl-CoA levels by catalyzing oxaloacetate decarboxylation. **a**, α-Keto acid and corresponding acyl-CoAs by decarboxylation and acylation. A hypothetical reaction of malonyl-CoA biosynthesis from oxaloacetate is indicated by brown box. **b**, Schematic of hypothetical metabolic networks of malonyl-CoA biosynthesis from oxaloacetate. **c-f**, Intracellular metabolite abundances of different strains or in different conditions. (**c**) Relative abundance of malonyl-CoA for the Control (*Escherichia coli* BW25113 strain), KDH, and KDH-ppc strains.Relative abundances of intracellular acetyl-CoA (**d**), oxaloacetate (**e**), and malonyl-CoA (**f**) for strain KDH treated with oxaloacetate or glucose. **g-i**, β-Alanine auxotrophy-dependent strain growth and the function of α-keto acid dehydrogenase BkdFDH. **g**, Endogenic biosynthetic pathway (red) for β-alanine and its complementary pathway (purple). **h**, Auxotrophic strain identification supplied with 0.1 g/L β-alanine in M9-glucose medium. **i**, Defective and complementary strain growth. ΔpanD: *panD* and *fadR* gene knockout; ΔpanD-mcrC: strain ΔpanD with the β-alanine biosynthetic pathway generated by overexpressing mcrC (malonyl-CoA reductase); ΔpanD-KDH: strain ΔpanD overexpressing bkdFDH on the chromosome *fadR* loci. ΔpanD-mcrC-KDH: strain ΔpanD-KDH with the β-alanine biosynthetic pathway.

We further carried out *in vivo* experiments using β-alanine auxotrophy-dependent strains to validate the LC-MS/MS data obtained above. By deleting the L-aspartate decarboxylase gene *panD* and overexpression of malonyl-CoA reductase C-domian gene *mcrC*, the cell growth of strains is coupled with intracellular malonyl-CoA level (Fig. 2g) ^20, 21^. In a ΔpanD background, the strain was unable growth on M9-glucose medium (MgSO_4_ and CaCl_2_ added) unless 0.1 g/L β-alanine was added (Fig. 2h). Strain ΔpanD-mcrC with the β-alanine biosynthetic pathway can grow slowly in M9-glucose medium (Fig. 2i), while the growth of ΔpanD-mcrC-KDH was almost fully recovered, indicating that the metabolic flux from glucose to malonyl-CoA is enhanced. The most plausible explanation for this result is that the increased malonyl-CoA level upon overexpressing the α-keto acid dehydrogenase complex provides more β-alanine for cell growth.

### A novel oxlaocetate dehydrogenation reaction cataylized by α-keto acid dehydrogenase

LC-MS/MS data combined with β-alanine auxotrophy-dependent strain growth experiments confirmed that expression of the gene cluster *bkdF-bkdG-bkdH-lpdA1* significantly increases the malonyl-CoA level, but the BkdFGH complex could not be distinguished from any of its components in terms of the cause of the results, since the E1 component can catalyze the α-keto acid decarboxylic reaction separately ^22^, and the E2 or E3 component of BkdFGH may assemble with the endogenous α-keto acid dehydrogenase complex ^23^. Subsequently, we deleted the subunits of this complex from strain ΔpanD-mcrC-KDH individually, obtaining strains ΔE1α (Δ*bkdF*), ΔE1β (Δ*bkdG*), ΔE2 (Δ*bkdH*), and Δlpd (Δ*lpdA1*). None of the strains with these complex subunit deficiencies exhibited cell growth recovery compared with strains ΔpanD-mcrC and ΔpanD-mcrC-KDH (Fig. 3a). Therefore, the results suggest that the entire α-keto acid dehydrogenase complex catalyzes the malonyl-CoA-producing reaction.

**Fig. 3.**
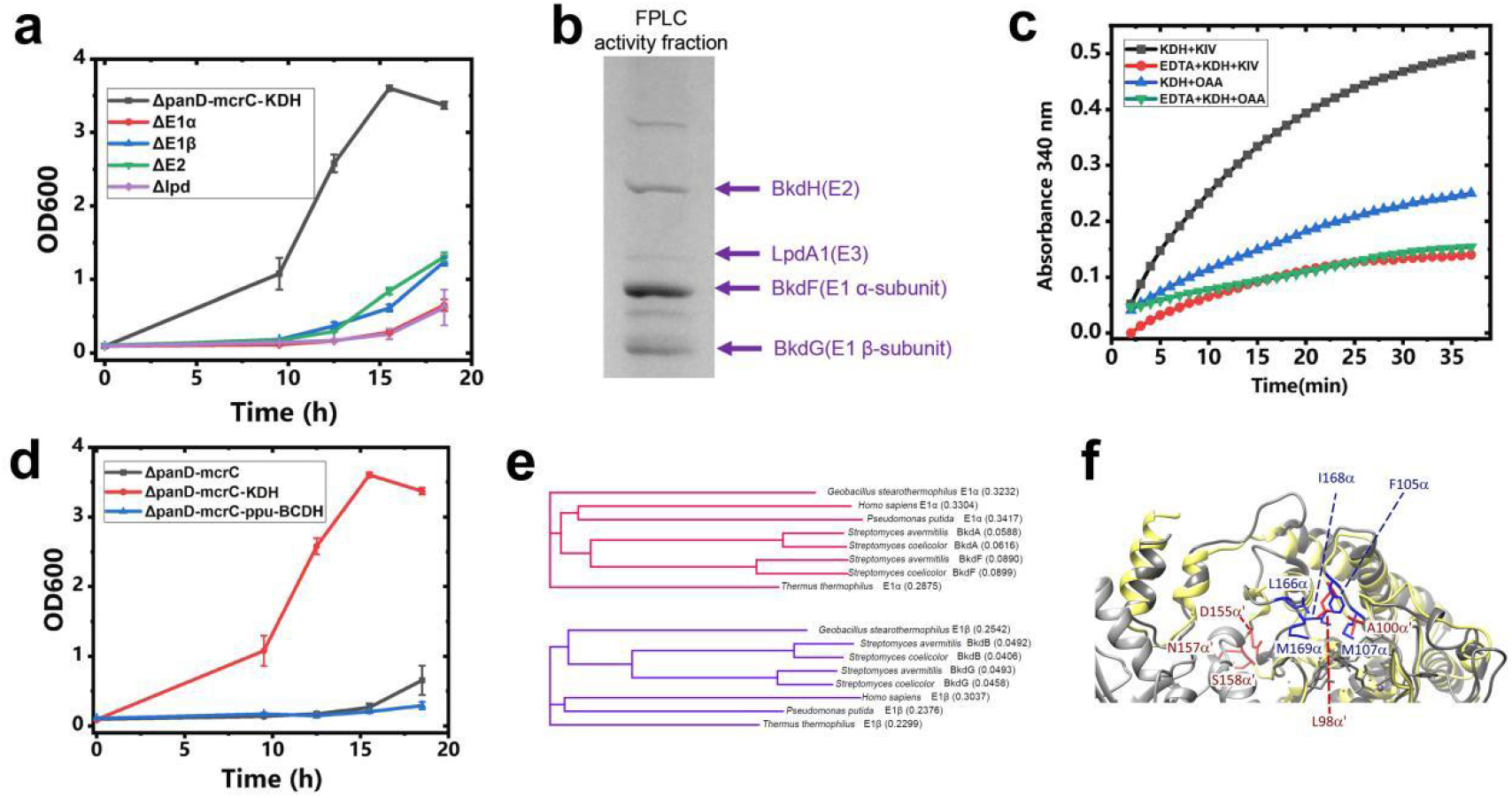
α-Keto acid dehydrogenase complex increases intracellular malonyl-CoA levels by catalyzing oxaloacetate decarboxylation. **a**, β-Alanine auxotrophy-dependent strain growth of different strains with subunit deficiency of BkdFDH. Strains ΔE1α, ΔE1β, ΔE2, and Δlpd were modified based on strain ΔpanD-mcrC-KDH with *bkdF, bkdG, bkdH* and *lpdA1* gene deletion, respectively. **b-c**, Purification and activity evaluation of the α-chain keto acid dehydrogenase complex. **b**, SDS-PAGE results for the α-chain keto acid dehydrogenase BkdFGH purified using FPLC and a 100-kDa cutoff Biomax ultrafiltration device; **c**, α-chain keto acid dehydrogenase activity of the reconstituted complex with α-keto-isovalerate and oxaloacetate. KDH+KIV: activity of substrate α-keto-isovalerate; KDH+OAA: activity of substrate oxaloacetate; EDTA+KDH+KIV and EDTA+KDH+OAA: Mg^2+^ was chelated by excess EDTA in the same test. **d**. Branched-chain α-keto acid dehydrogenase from *Pseudomonas putida* does not promote β-alanine auxotrophy-dependent strain cell growth. Strain ΔpanD-mcrC-ppu-BCDH: β-alanine auxotrophy-dependent strain with a malonyl-CoA replenishment pathway and overexpressed *Pseudomonas putida* branched-chain α-keto acid dehydrogenase complex. **e**. Sequence comparison of *S. avermitilis* BkdF/BkdG with other E1α/E1β of *S. avermitilis* BkdABC, *Streptomyce*s *coelicolor* BkdFGH and BkdABC, and 4 structure-solved branched-chain α-keto acid dehydrogenase from other species. **f**. Comparison of a predicted structure of *S. avermitilis* BkdF using SWISS-MODEL homology modeling and *P. putida* branched-chain α-keto acid dehydrogenase complex E1α/E1β. Structure of *P. putida* E1α/E1β are shown in gray, *S. avermitilis* BkdF is shown in yellow. The conserved residues for substrate-binding in *P. putida* E1α and corresponding residues in *S. avermitilis* BkdF are shown in blue and red, respectively.

To further verify this novel reaction, we purified this enzyme (see supplementary S1) and tested its activity on different substrates (Fig. 3b). The purified α-chain keto acid dehydrogenase complex BkdFGH was found to be active with both α-keto-isovalerate (specific activity 9.677 ± 0.130 μmol/min/mg) and oxaloacetate (specific activity 4.705 ± 0.617 μmol/min/mg). The E1 component catalyzes the first decarboxylation and acylation of α-keto acids, and it also participates in substrate recognition ^24^. The key activator of the E1 component, Mg^2+ 25^, is chelated by excess EDTA, resulting in almost complete activity loss, both with α-keto-isovalerate and oxaloacetate (Fig. 3c). The data obtained here combined with stable isotopic labeling metabolic flux analysis results (see Supplementary Fig. S2) confirmed that α-chain keto acid dehydrogenase BkdFGH from *S. avermitilis* (MA-4680) has a broad substrate spectrum (α-keto-isovalerate, α-keto-β-methyl-valerate, and α-keto-γ-methyl-valerate) that includes oxaloacetate, as proposed here.

In particular, we have confirmed for the first time that malonyl-CoA is generated by oxaloacetate decarboxylation.

It is of interesting to investigate if the branched-chain α-keto acid dehydrogenases from other species also has oxaloacetate dehydrogenase activity. To test this hypothesis, we overexpressed the well-studied branched-chain α-keto acid dehydrogenase complex (Accession: M57613) from *Pseudomonas putida* by integrating the whole expression cassette into the *fadR* loci of strain RD, obtaining strain RD-ppu-BCDH. The pS95s-MCR-C plasmid expressing malonyl-CoA reductase (Supplementary Table S1) was then transformed into strain RD-ppu-BCDH, generating strain ΔpanD-mcrC-ppu-BCDH. We then compared the cell growth of this strain and that of ΔpanD-mcrC-BCDH in M9-glucose medium. Unexpectedly, unlike the complex from *S. avermitilis*, the branched-chain α-keto acid dehydrogenase complex from *Pseudomonas putida* does not promote β-alanine auxotrophy-dependent strain growth (Fig. 3d), suggesting that the former does not increase intracellular malonyl-CoA levels and oxaloacetate dehydrogenation is not a comprehensive non-specific reaction of branched-chain α-keto acid dehydrogenases. The molecular mechanism of oxaloacetate substrate recognition by *S. avermitilis* BkdFGH is still to be further explored. However, according to our structural modelling and sequence comparisons, *S. avermitilis* BkdFGH might differ from other branched-chain α-keto acid dehydrogenases at both structure and sequence levels, but is very close to BkdFGH from other *Streptomyce*s species, such as *Streptomyce*s *coelicolor* (Fig.3e-f).

### Oxaloacetate dehydrogenase (OADH) plays significant role in the regulation of malonyl-CoA-derived chemical production in its native host

Previous studies on the branched-chain α-keto acid dehydrogenase complex from *Streptomyces* mainly focus on branched-chain amino acid degradation and corresponding branched-chain acyl-CoA production for secondary metabolite biosynthesis ^19^. However, in the present study, we have confirmed that the branched-chain α-keto acid dehydrogenase complex BkdFGH catalyzes decarboxylation of oxaloacetate to form malonyl-CoA. We suspected this enzyme complex also has important physiological significance by suppling malonyl-CoA ^26, 27^.

*S. avermitilis* natively produces avermectins (B1a) via condensation of one molecule of 2-methylbutanoyl-CoA, five molecules of methylmalonyl-CoA, and seven molecules of malonyl-CoA, where 2-methylbutanoyl-CoA and methylmalonyl-CoA are produced by dehydrogenation reactions catalyzed by BkdFGH ^28^. Clearly, *S. avermitilis* requires more malonyl-CoA than *P. putida*, which natively produces fatty acids instead of polyketides or other malonyl-CoA derivatives for growth.

*S. coelicolor* M145 is a close relative of *S. avermitilis* that natively produces actinorhodin by condensing 16 molecules of malonyl-CoA ^29^. *S. coelicolor* M145 contains two sets of branched-chain α-keto acid dehydrogenase complex genes (named *bkdABC* and *bkdFGH*), much like *S. avermitilis* ^30^. However, unlike *S. avermitilis*, the natural secondary product of *S. coelicolor* M145 is synthesized from a single precursor (Fig. 4a). Therefore, it is an ideal model organism for investigating the relationship between the branched-chain α-keto acid dehydrogenase complex and intracellular malonyl-CoA production.

**Fig. 4.**
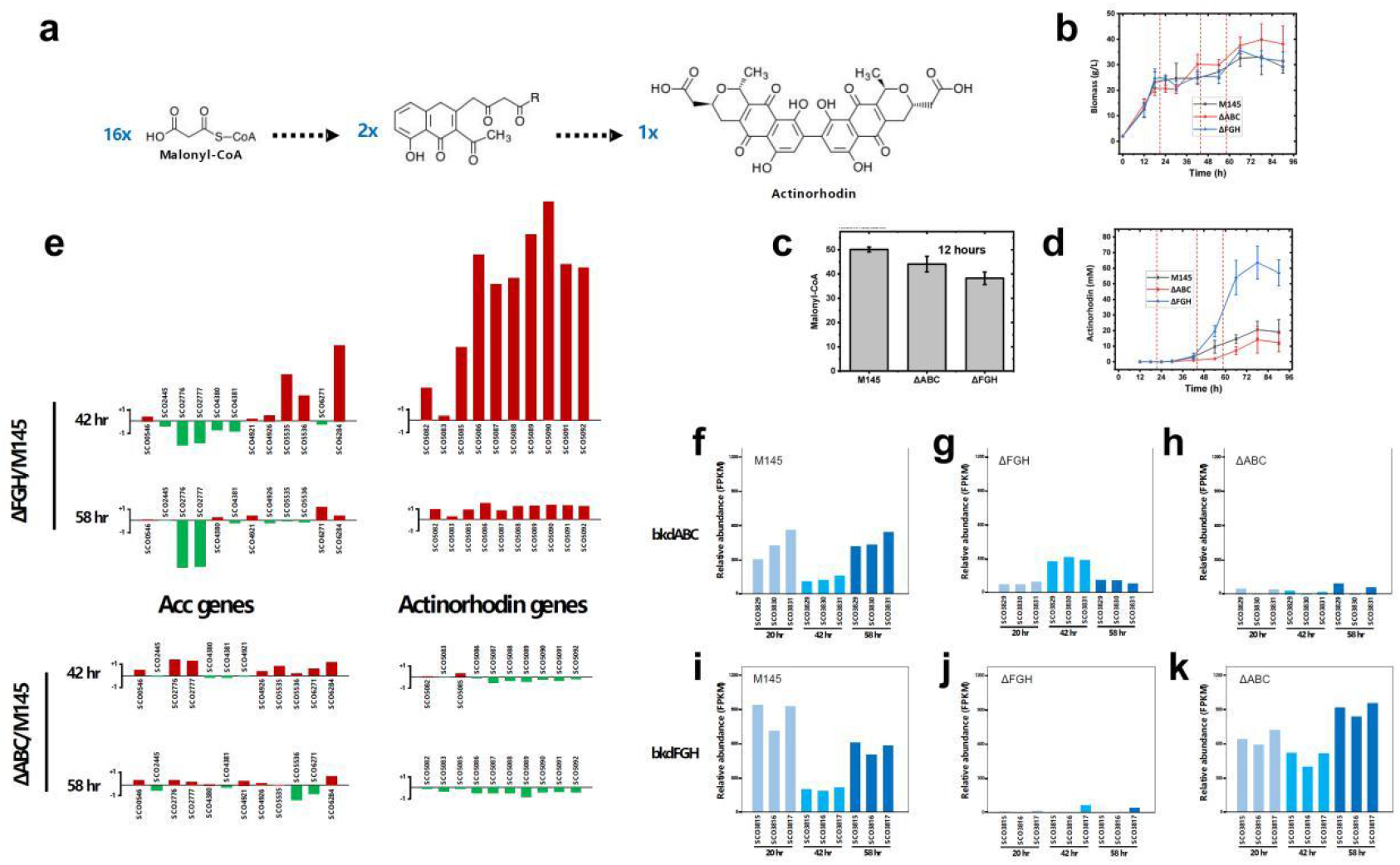
Physiological function of branched-chain α-keto acid dehydrogenase in *Streptomyces*. **a**, Actinorhodin biosynthesis in *Streptomyces coelicolor* via condensation of 16 molecules of malonyl-CoA. **b**, Growth curves for strains M145, ΔABC, and ΔFGH. M145: wild-type strain of *S. coelicolor* M145; ΔABC: gene *bkdABC* knockout mutant; ΔFGH: gene *bkdFGH* knockout mutant. **c**, Relative intracellular malonyl-CoA abundances for strains M145, ΔABC, and ΔFGH. **d**, Actinorhodin production for strains M145, ΔABC, and ΔFGH. **e-k**, Transcriptome analysis of α-keto acid dehydrogenase gene mutant strains. **e**, Differential expression of Acc and Actinorhodin genes for mutant strains as compared with those of wild-type M145. Acc: acetyl-CoA carboxylase; Actinorhodin: biosynthesis gene cluster (for gene annotations see Supplementary Information). **f-k**, Relative branched-chain α-keto acid dehydrogenase gene expression levels for mutant strains in different growth phases. SCO3831: *bkdA*, E1 alpha subunit; SCO3830: *bkdB*, E1 beta subunit; SCO3829: *bkdC*, E2 core; SCO3817: *bkdF*, E1 alpha subunit; SCO3816: *bkdG*, E1 beta subunit; SCO3815: *bkdH*, E2 core.

In the present study, we knocked out the *bkdABC* and *bkdFGH* genes of *S. coelicolor* M145 (mutants were named ΔABC and ΔFGH, respectively) to determine whether the α-keto acid dehydrogenase complex is necessary for cell growth or only for the secondary-metabolite-accumulation period (Fig.4b-d). *bkdABC* and *bkdFGH* knockout reduces intracellular malonyl-CoA levels by 12.21% and 24.03% in the mid-log phase (12 h, Fig. 4c), respectively, showing that malonyl-CoA production is increased by oxaloacetate dehydrogenase in the cell-growth period, in which *bkdFGH* is dominant.

It was shown *bkdABC* knockout reduces actinorhodin production by 49.72% in the stationary phase (66 h) (Fig. 4d), while *bkdFGH* knockout increases actinorhodin production 2.81-fold in the same phase. These puzzling results left us confused as to what processes promote actinorhodin production inside the cell. Accordingly, we performed an integrated transcriptome analysis of *S. coelicolor* M145 and its mutants. Acetyl-CoA carboxylase, actinorhodin biosynthesis, and branched-chain α-keto acid dehydrogenase genes were studied in particular (Fig. 4e-k). The expression levels of acetyl/propionyl-CoA-carboxylase-related genes are presented in Fig. 4e. Compared with the wild-type strain (M145), the expression levels of both Acc and Actinorhodin biosynthesis genes for the *bkdABC* knockout strain (ΔABC) are not significantly different in the mid-to late-stationary phase (42 and 58 h), while those for the *bkdFGH* knockout strain (ΔFGH) are clearly up- or downregulated. The acetyl-CoA-carboxylase-related genes SCO5535, SCO5536, and SCO6284 are upregulated 3.78-, 2.08-, and 6.22-fold in the mid-stationary phase (42 h) when actinorhodin synthesis by the ΔFGH strain starts, among which SCO5535 (*accB*) and SCO5536 (*accE*) are key acyl-CoA carboxylase components ^31^, and SCO6284 is an acyl-CoA carboxylase beta subunit.

We speculate that the acetyl-CoA carboxylase genes of strain ΔFGH are compensatorily upregulated in the mid-stationary phase (42 h), and the consequently increased malonyl-CoA triggers actinorhodin biosynthesis, resulting in an increase in actinorhodin production. However, this hypothesis requires more evidence. The *bkdABC* and *bkdFGH* expression levels in the wild-type strain are high in the cell-growth (20 h) and actinorhodin-accumulation periods (58 h) (, Fig. 4f-k). Thus, the results presented here show that α-keto acid dehydrogenase in *Streptomyces* is required in both the cell-growth and secondary metabolite accumulation periods, during which malonyl-CoA is used for fatty acid and polyketide biosynthesis. And oxaloacetate dehydrogenases indeed involved in the regulation of malonyl-CoA-derived actinorhodin biosynthesis.

### Supply of malonyl-CoA from oxaloacetate by OADH is a novel and efficient biotechnological route for the production of malonyl-CoA-derived chemicals

The novel malonyl-CoA biosynthesis pathway has significant biotechnological application potential in the production of malonyl-CoA-derivative chemicals. The decarboxylation, energy-release, and irreversible reactions catalyzed by OADH have thermodynamic and effiicency advantages over the ATP-consuming and CO_2_-fixation reactions catalyzed by acetyl-CoA carboxylase (Fig. 5c). Accordingly, by engineering OADH module, we developed several engineering strains to efficiently produce malonyl-CoA derivatives, including 3-hydroxypropionate, triacetate lactone, phloroglucinol, aloesone, flaviolin, raspberry ketone, and resveratrol (Fig. 5a-b).

**Fig. 5.**
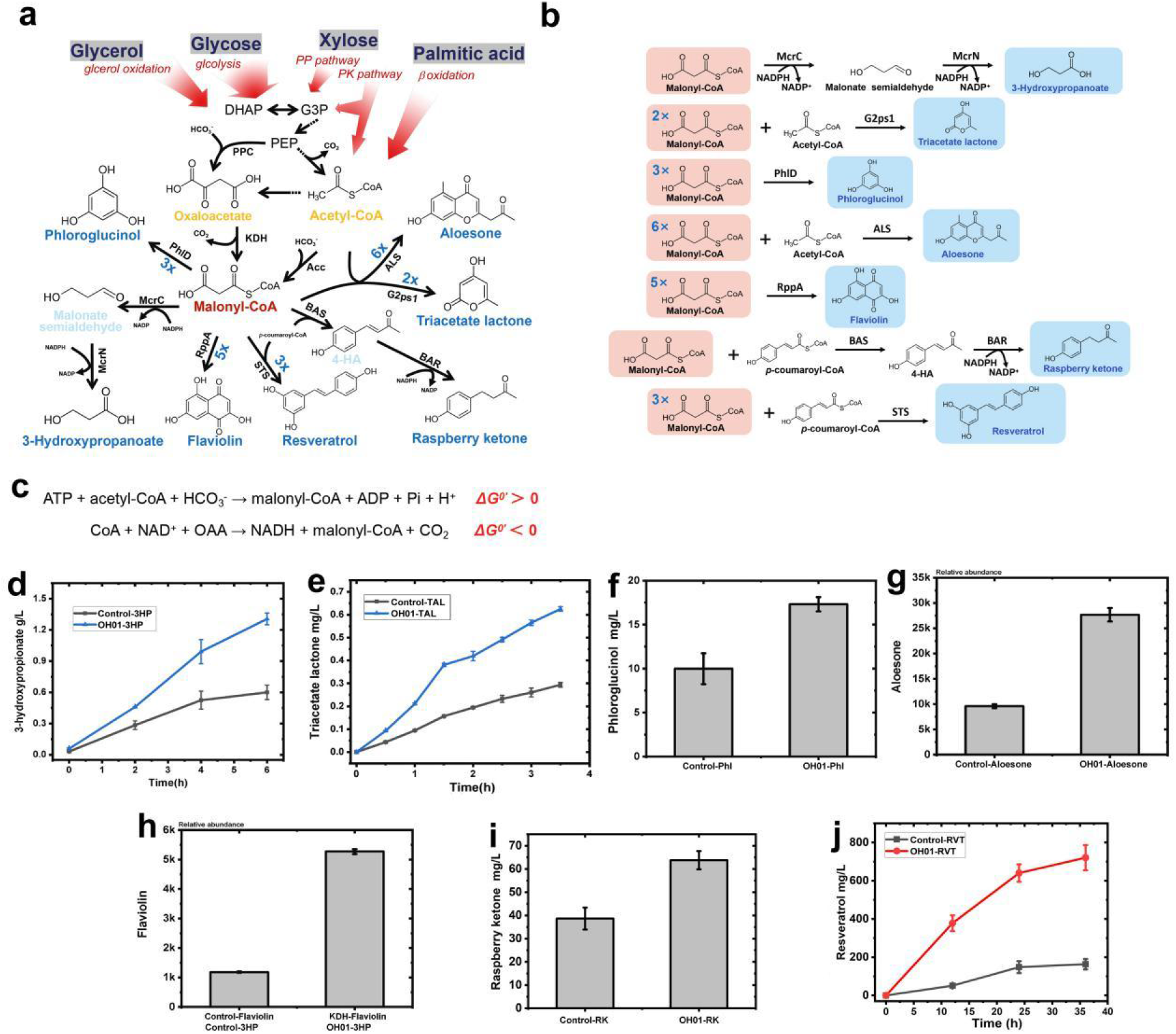
Utility evaluation of the novel malonyl-CoA-derivative biosynthesis platform. **a-b**, Novel biosynthetic pathways to 3-hydroxypropionate, triacetate lactone, phloroglucinol, aloesone, and flaviolin. Acetyl-CoA comes from glucose (thourgh glycolysis), glycerol (through glycerol oxidation), xylose (through pentose phosphate pathway (PP pathway) or phosphoketolase pathway (PK pathway), or palmitic acid (through β-oxidation). McrC and McrN: malonyl-CoA reductase from *Chloroflexus aurantiacus*; G2ps1: 2-pyrone synthase from *Gerbera hybrid*; ALS: aloesone synthase from *Rheum palmatum*; PhlD: 1,3,5-trihydroxy synthase from *Pseudomonas protegens*; RppA: 1,3,6,8-tetrahydroxynaphthalene synthase from *Streptomyces griseus*. **c**, Comparison of biosynthesis of malonyl-CoA through the two different reaction. **d-e**, Production and productivity comparison of 3-hydroxypropionate and triacetate lactone. **f-i**, Production comparison of phloroglucinol, aloesone, flaviolin, and raspberry ketone after 0.5-, 3-, 3-,24h bioconversion, respectively. **j**, Production comparison of resveratrol.

3-Hydroxypropionate, an important platform chemical, was produced using malonyl-CoA as the only precursor via two reduction steps catalyzed by malonyl-CoA reductase ^20^. The 3-hydroxypropionate biosynthesis module (pSB1s-MCR-CN) was introduced into OH01 strain in which OADH module was enhanced and optimized (resulting OH01-3HP). It was shown the titer and productivity of 3-hydroxypropionate can be increased by 3-folds by this novel malonyl-CoA platform, especially in the first 4 h (Fig.5d).

Triacetate lactone (TAL), a typical malonyl-CoA-derivative chemical, can be formed using an acetyl-CoA as starter unit and two malonyl-CoA as extender units by type III PKS 2-pyrone synthase from *Gerbera hybrid* encoded by g2ps1 (Accession: P48391) ^32^. TAL biosynthesis module was transformed into OH01 strain (resulting OH01-TAL). As expected, OADH module significantly promotes TAL titer and productivity by 3-folds, especially in the first 2 h (Fig.5e).

Aloesone is an aromatic heptaketide produced from one molecule of acetyl-CoA and six molecules of malonyl-CoA naturally in *Rheum palmatum* ^33^. It was employed here to evaluate the ability of strain OH01 to supply malonyl-CoA unit. The engineered strain OH01-Aloesone was obtained by expressing aloesone synthase (Accession: AAS87170). Similar to above results, enhancement of OADH module also can significantly improve aloesone production. After 3 h of bioconversion, KDH-Aloesone shows production approximately 3-folds that of Control-Aloesone (Fig. 5f).

Phloroglucinol and flaviolin are polyketides that require malonyl-CoA as both the starter and extender unit, condensing three and five malonyl-CoA units, respectively (Fig. 5b). The phlD gene coding 1,3,5-trihydroxy synthase (Accession: WP_011064130.1) from *Pseudomonas protegens* and the rppA gene coding 1,3,6,8-tetrahydroxynaphthalene synthase (Accession: AXN70078) from *Streptomyces griseus* ^34^ were cloned into plasmid pXB1k and pLB1s and expressed in the control and OH01 strains, respectively. In the first 30 min, a 100% increase in phloroglucinol production was achieved using the oxaloacetate dehydrogenase complex (Fig. 5g). Analogously, overexpressing the oxaloacetic acid dehydrogenase complex increases flaviolin production 4.5-fold in 3 h (Fig. 5h).

Phenylpropanoids, including flavonoids, are a class of polyphenolic secondary metabolites and widely used in pharmaceuticals and cosmetics. They are synthesized by the phenylpropanoid metabolic pathway, and chalcone synthase (CHS) catalyses the key step using malonyl-CoA and phenylalanine-derived *p*-coumaroyl-CoA as precursors. Raspberry ketone and resveratrol were used as products to illustrate the effect of OADH module during phenylpropanoids biosynthesis. Glycerol was used as carbon source and coumaric acid was added in medium. It was shown when raspberry ketone biosynthesis module consists of BAS, BAR, and 4CL was introduced into OH01 strain, raspberry ketone production could increase by 50% (Fig. 5i). As for resveratrol production, we introduced and optimized resveratol biosynthesis module into OH01 strain, and the resulting OH01-RVT strain could produce about 720.54 mg/L resveratol, about 4.4-folds of control strain (Fig. 5j).

Taken together, we have confirmed significant application potential of OADH in the biosynthesis of malonyl-CoA derivatives. Production of the non-polyketide compound 3-hydroxypropionate; the multi-precursor polyketides triacetate lactone and aloesone; sole-precursor polyketides phloroglucinol and flaviolin; phenylpropanoids raspberry ketone; and flavonoids resveratrol, were all observably promoted by OADH. Therefore, this novel bio-manufacturing platform can be readily employed in the biosynthesis of malonyl-CoA derivatives.

## Discussion

ΔThe physiological significance of oxaloacetate dehydrogenase in its native hosts is another focus of future studies. Indeed, in literatures 2-4 decades ago, it has postulated there is oxaloacetate dehydrogenase that could catalyze generation of malonyl-CoA from oxaloacetate in *Streptomyces aureofuciens* ^37, 38^. However, further study did not supported these claims and no genes of oxaloacetate dehydrogenases have been identified in any species yet ^39^. Eventually, in this study, we got it from *Streptomyces* and resolved a long-time conjecture, and therefore expand the knowledge of enzyme species.

Some results of our study still need to be further investigated. The knocked-out of *bkdFGH* in *S. coelicolor* indeed lead to reduce of malonyl-CoA level. However, the production of malonyl-CoA-derived actinorhodin, on the contrary, increased significantly. This result was not surprising as there are several components of acetyl-CoA carboxylase or oxaloacetate dehydrogenases in *S. coelicolor*. A global regulation between different malonyl-CoA-producing enzymes and actinorhodin biosynthesis pathway might occur in given conditions. Several results of previous works were consistent with our results. For example, increased phosphoenolpyruvate carboxylase (PPC) activity was observed during actinorhodin biosynthesis in *S. coelicolor* ^40^; nitrogen conditions can affect the polyketide biosynthesis and potential malonyl-CoA biosynthesis enzymes activity in *S. aureofuciens* ^38^. These results combined with our study implied oxaloacetate dehydrogenase might play an important role in the regulation between primary and secondary metabolism.

Furthermore, we argued the real physiological roles of BkdFGH in *S. avermitilis* in previous reports ^19^. It is long believed that BkdFGH is responsible for the biosynthesis of the starter unit of avermectin in *S. avermitilis* and thus significantly affect the production of avermectin. In fact, there are no adequate metabolic and biochemical evidences regarding the function of *S. avermitilis* BkdFGH. Therefore, it is necessary to carry out research to further elucidate its physiological roles. Our work presents a novel and effective strategy for increasing intracellular malonyl-CoA levels that can be used in the biosynthesis of numerous value-added malonyl-CoA derivatives. Biosynthesis of malonyl-CoA through PEP-oxaloacetate pathway has a carbon-fixation reaction catalyzed by phosphoenolpyruvate carboxylase and a carbon-loss reaction catalyzed by oxaloacetate dehydrogenase, while through pyruvate-acetyl-CoA pathway has a carbon-loss step catalyzed by pyruvate dehydrogenase and then a carbon-fixation step catalyzed by acetyl-CoA carboxylase. Because the carboxylation efficiency of phosphoenolpyruvate carboxylase is higher than that of acetyl-CoA carboxylase ^7^, the former pathway might be a more efficient choice for the supply of malonyl-CoA in biotechnological fields.

Furthermore, the unique acetyl-CoA carboxylation strategy for malonyl-CoA supply has its drawbacks: for example, improvement of acetyl-CoA pool might lead to increased acetate accumulation; modification of acetyl-CoA utilization pathway might reduce TCA cycle activity thus influence the energy metabolism and cell growth. The additional malonyl-CoA-producing pathway will be helpful for development of more flexible metabolic engineering strategies. Oxalocetate can be supply from diverse carbon sources, such as glucose, glycerol, xylose, and fatty acids. This will facilitate to develop different carbon-source strategies favorable both chemicals production and cell growth. Take together, we have provided a new way for malonyl-CoA biosynthesis as well as a novel biotechnological route for production of malonyl-CoA derivatives chemicals.

## Methods

### Strains and reagents

The *E. coli* strains DH5α and BW25113/F′ were used as host strains for plasmid construction. The *E. coli* strain BW25113/F′ and *S. coelicolor* strain M145 were used for related tests. Chemicals were from Sigma-Aldrich (St. Louis, MO, USA) unless otherwise specified. Fast Pfu Fly™ DNA polymerase was obtained from Trans Gen Biotech Co., Ltd. (Beijing, China). A Gibson Assembly® Cloning Kit (New England Biolabs, Beverly, MA, USA) was used for plasmid construction. All strains and plasmids used in this study are listed in Table S1 and detailed information is provided.

Luria-Bertani (LB) medium (10 g/L tryptone, 5 g/L yeast extract, and 10 g/L NaCl) and ZYM medium (10 g/L tryptone, 5 g/L yeast extract, 25 mM Na_2_HPO_4_, 25 mM KH_2_PO_4_, 50 mM NH_4_Cl, 5 mM Na_2_SO_4_, 5 g/L glycerol, 0.5 g/L glucose, 2 g/L L-arabinose) were used to grow *E. coli* cells and express proteins, respectively. When necessary, antibiotics or MgSO_4_ were added (to a final concentration of 100 mg/L for carbenicillin, 50 mg/L for kanamycin, 50 mg/L for streptomycin, and 2 mM for MgSO_4_). The *S. coelicolor* M145 strain and the BCDH mutant strains were cultured in YEME medium (3 g/L malt extract, 3 g/L yeast extract, 5 g/L peptone, 10 g/L glucose, 103 g/L sucrose and 5 mM MgCl_2_). R5 medium (103 g/L sucrose, 10 g/L glucose, 5 g/L yeast extract, 0.1g/L casamino acids, 10.12 g/L MgCl_2_·6H_2_O, 0.25 g/L K_2_SO_4_, 25 mM TES, 0.08 mg/L ZnCl_2_, 0.4 mg/L FeCl_3_·6H_2_O, 0.02 mg/L CuCl_2_·2H_2_O, 0.02/L mg MnCl_2_·4H_2_O, 0.02 mg/L Na_2_B_4_O_7_·10H_2_O, 0.02 mg/L (NH_4_)6Mo_7_O_24_·4H_2_O) was used for actinorhodin production.

The whole-cell bioconversion for intracellular metabolite quantitative analysis was carried out using M9 medium containing 12.8 g/L Na_2_HPO_4_·7H_2_O, 3 g/L KH_2_PO_4_, 0.5 g/L NaCl, and 1 g/L NH_4_Cl, and 10 g/L glucose or 3.67 g/L oxaloacetate. For ^13^C-labeling metabolic flux analysis, [U-^12^C_6_] glucose was replaced by stable isotope labeling glucose [U-^13^C_6_] and 150 mM NaH^12^CO_3_ was added. M9-glucose medium supplemented with 0.1 mM CaCl_2_·2H_2_O and 2 mM MgSO_4_ was used for deficient strain growth study. To produce 3-hydroxypropionate by whole-cell bioconversion, 2 g/L fumarate, 2 g/L oxaloacetate, and 3 g/L palmitic acid (emulsified with 0.6 g/L Brij 58) were added to the M9 medium. To produce triacetate lactone, phloroglucinol, aloesone, and flaviolin by whole-cell bioconversion, 10 g/L glucose and 4 g/L fumarate were added.

### DNA manipulation and genome editing

Molecular cloning was performed using standard protocols ^41^. *E. coli* gene-deletion strains were obtained from the Keio Collection ^42^. For *E. coli* genome integration of α-keto acid dehydrogenase BdkFGH and acetyl-CoA carboxylase (ACC) genes, the fragments of P_119_-*bkdFGH-lpdA1* or P_119_-*accBCDA* cassette, flanked with the Lox66-kan^R^-Lox71 cassette, were amplified from pSL91k-BCDH and pSL91k-ACC, respectively. Then, the gene-Lox66-kan^R^-Lox71 fragments were assembled with approximately 1000 nt of the up/downstream regions of the target genes by overlap-extension PCR, resulting in the target fragments. *E. coli* genome integration was carried out using λ Red recombineering as previously described ^43^. The *E. coli* kanamycin resistance marker was eliminated using the plasmid pSB1s*-Cre by adding 0.2% L-arabinose or by using pCP20. For *E. coli* chromosomal integration and gene deletion of multiple targets, P1 virus-mediated transfection was performed, as described elsewhere ^44^. Gene knockout of *S. coelicolor* M145 has been described previously ^45^. Mutant strains were verified by PCR and further confirmed by sequencing (GENEWIZ, Suzhou, China).

### Intracellular metabolite determination and protein verification

For intracellular metabolite determination, strains were grown at 37 °C in LB medium to the early-stationary phase. Cells were collected at 0–4 °C by centrifugation at 5,000 ×g for 10 min, washed twice with ice-cold 0.85% NaCl solution, and then resuspended in M9-glucose medium to OD_600_ = 30. After shaking at 220 rpm in tubes at 37 °C for 2.5 h, cells were recollected and washed as mentioned above, and then resuspended in 400 μL (OD_600_ = 250) 80% (*v/v*) aqueous methanol solvent (prechilled to -80 °C) immediately, sonicated, and concentrated at the critical point as described previously ^16^. Samples were redissolved in 50 μL ice-cold 80% (*v/v*) aqueous methanol and injected into an LC-MS/MS system for qualitative and quantitative analysis.

The LC-MS/MS system consisted of a Dionex Ultimate 3000 UPLC unit (Dionex, Sunnyvale, CA, USA) coupled to a TSQ Quantiva Ultra triple-quadrupole mass spectrometer (Thermo Fisher, Waltham, MA, USA) equipped with a heated electrospray ionization (HESI) probe. Detailed LC and MS conditions were described previously ^21^. Data analysis and quantification were performed using Xcalibur 3.0.63 software (Thermo Fisher). In particular, data for malonyl-CoA (m0), and its 1(m1), 2(m2), and 3(m3) carbon isotope labels were acquired in selected reaction monitoring (SRM) with transitions of 854/347, 855/348, 856/349, and 857/350, respectively. Both precursor and fragment ions were collected with a resolution of 0.7 FWHM, respectively. The source parameters are as follows: spray voltage: 3500 V; ion transfer tube temperature: 350 °C; vaporizer temperature: 300 °C; sheath gas flow rate: 30 Arb; auxiliary gas flow rate: 10 Arb. CID gas: 1.5 m Torr.

For protein verification, the cut gel pieces containing the target proteins were vacuum-dried, destained, and digested with 12.5 ng/μL trypsin in 50 mM ammonium bicarbonate (pH 8.0) for 16 h at 37 °C. Then, the digested supernatants were mixed with the MALDI matrix material (5.0 mg/mL of α-cyano-4-hydroxycinnamic acid) at 1:1 ratio and spotted onto a MALDI target. After air drying, crystallized spots were analyzed with a MALDI-TOF/TOF analyzer (AB SCIEX 5800) (AB Sciex, CA, USA) in positive reflection mode. Parent mass peaks in the mass range 800–3500 Da and the ten most abundant ions from MS analysis were chosen for tandem TOF/TOF analysis.

The mass spectra were analyzed using the MASCOT search engine (Matrix Science, London, UK) to search against the *E. coli* or *S. avermitilis* database. The search parameters were as follows: two missed cleavage sites by trypsin, 0.2 Da peptide mass tolerance, 0.5 Da MS/MS ion tolerance, carbamidomethylation of cysteine as a fixed modification, and methionine oxidation as a variable modification.

### Determination of cell growth

*E. coli* strain growth was monitored by measuring changes in OD_600_. Tested strains were cultured overnight in LB medium, washed with 0.85% NaCl, suspended in 50 μL (OD_600_ = 5) 0.85% NaCl solution, and sub-inoculated (1:100) into 5 mL M9-glucose medium (37 °C, 220 rpm). A 200-μL aliquot of the culture was sampled every time and then measured using a UV-2550 spectrophotometer (Shimadzu Corp., Kyoto, Japan). To measure *S. coelicolor* M145 biomass, 1 mL fermentation broth was centrifugated and the pellets were washed twice with distilled water to determine the fresh weight.

### Purification and activity analysis of BkdFGH

Proteins were fused with His_6_-Tag for Ni^2+^-NTA affinity purification as previously described with some modifications ^22, 46^. Strains were cultured in LB medium containing 2 g/L L-valine and 2 mM MgSO_4_·7H_2_O at 30 °C for 20 h. Cells were harvested at 0–4 °C by centrifugation at 5,000 × g for 20 min, washed twice with ice-cold buffer A (50 mM Tris-HCl, 200 mM KCl, pH 8.0 at 4 °C, filtered through a 0.22-μM Millipore membrane), resuspended in buffer A with 20 mM imidazole added, and then disrupted using an AH-BASIC high-pressure homogenizer (ATS Engineering Inc., Canada). The cell disruption suspension was centrifuged at 10,000 × g for 1 h three times and the supernatant was filtered through a 0.22-μM Millipore membrane to further remove as much precipitate as possible before being packed into a column.

His_6_-protein was eluted in sequence with 20-, 40-, 80-, and 250-mM imidazole in buffer A. Column fraction (250 mM imidazole) was filtered using an ultrafiltration device (100-kDa cutoff Biomax for BCDH-His_6_, 30-kDa cutoff Biomax for LpdA1-His_6_) at 0–4 °C and 8,000 × g and washed thrice with buffer B (buffer A with 0.2 mM DTT added) to remove imidazole and small molecular weight impurity proteins. LpdA1-His_6_ was stored at -80 °C in enzyme storage buffer C (buffer B with 20% glycerol, 0.5 mM ThDP, and 1 mM MgCl_2_). For further purification of BCDH-His_6_, the concentrated protein was applied to an FPLC (AKTA) Superose 6 Increase 10/300 GL gel-filtration column equilibrated with buffer A (all FPLC conditions given in Supplementary Fig. S1). The enzyme was re-concentrated using the ultrafiltration device mentioned above and stored at -80 °C in buffer C.

Assay buffer D composed of 100 mM Tris-HCl (pH 7.2 at 30 °C), 2 mM MgC1_2_, 1 mM ThDP, 1 mM CoA, 6 mM NAD^+^, and 0.4 mM DTT was prepared freshly on ice and prewarmed to 30 °C for the following spectrophotometric analysis. The total volume of the assay mixture for the spectrophotometric assay was 200 μL, which comprised 100 μL buffer D and 90 μL ddH_2_O prewarmed to 30 °C in 96-well plate, then 5 μL of the entire complex assembled above was added to the assay buffer and mixed thoroughly. The reaction is initiated with 5 μL of 160 mM α-keto-isovalerate or oxaloacetate (dissolved in 100 mM Tris-HCl, pH 7.2 at 30 °C) prewarmed to 30 °C and the absorbance at 340 nm was measured immediately using a microplate reader (SynergyMx, BioTek, VT, USA) for 1 h. When necessary, 10 mM EDTA was added to the assay buffer to chelate Mg^2+^, the catalytic center ion of the E1 component ^47^.

### Production of 3-hydroxypropionate, triacetate lactone, phloroglucinol, aloesone, flaviolin, raspberry ketone, resveratrol, and actinorhodin

Test strains were induced to the stationary phase at 37 °C in ZYM medium, harvested by centrifugation at 5,000 × g and 0–4 °C, and resuspended in the bioconversion buffer mentioned above to OD_600_ = 30. The 3-hydroxypropionate, triacetate lactone, phloroglucinol, raspberry ketone, and resveratrol concentrations were then determined using an HPLC apparatus (LC-20A LabSolutions, Shimadzu Corp., Kyoto, Japan) equipped with a photo-diode array and a refractive index (RI) detector. 3-Hydroxypropionate samples were eluted through a 300 mm × 7.8 mm Aminex HPX-87H column (Bio-Rad, Hercules, CA, USA) at 55 °C using 5 mM H_2_SO_4_ (flow rate 0.6 mL/min) in 20 min. Triacetate lactone and phloroglucinol samples were analyzed with a 150 mm × 4.6 mm Shim-pack GIST C18-AQ column at 40 °C, eluted with 1 g/L formic acid:methanol (80:20 and 95:5), and detected at 283 and 254 nm, respectively. Relative abundances of aloesone and flaviolin were determined by LC-MS/MS. Detailed methods for aloesone and flaviolin production are described elsewhere ^2^.Raspberry ketone was anaylized with 250 mm×4.6 mm Agilent Extend-C18 column at 35 °C with a flow rate of 0.5 ml/min. Mobile phase A (65%) was water with 0.1% (v/v) formic acid; mobile phase B (35%) was acetonitrile. Raspberry ketone was detected via DAD detection at 222 nm. Resveratrol was analyzed with 250 mm × 4.6 cm Licrosphere RP C18 reverse-phase column with a flow rate of 0.5 ml/min. Mobile phases used were 10 % (v/v) MeOH in water (solvent A) and 10 % (v/v) water in MeOH (solvent B), resveratrol was detected via DAD at 280 and 306 nm.

For the measurement of actinorhodin production, 1 mL fermentation broth was extracted with 5 M KOH (at final 1 M concentration) and then shaken at 220 rpm for 15 min at room temperature. Supernatants were acquired by centrifugation at 4 °C, 13,000 rpm for 10 min and actinorhodin was quantified by UV absorbance at 640 nm using a Synergy H4 spectrophotometer (BioTek, USA) (Ɛ_640_ = 25,320).

### Transcriptome sequencing and analysis

RNA extraction was conducted as described previously ^48^. A total amount of 3 μg RNA per sample was used as the input material for RNA sample preparation. For prokaryotic samples, mRNA was purified from total RNA using probes to remove the rRNA. In order to select cDNA fragments preferentially of 370–420 bp in length, the library fragments were purified with an AMPure XP system (Beckman Coulter, Beverly, USA). Clustering of the index-coded samples was performed on a cBot Cluster Generation System using TruSeq PE Cluster Kit v3-cBot-HS (Illumia) according to the manufacturer’s instructions. After cluster generation, the library preparations were sequenced on an Illumina Novaseq platform and 150 bp paired-end reads were generated.

Differential expression analysis between two conditions (three biological replicates per condition) was performed using the DESeq R software package (1.18.0). DESeq provides statistical routines for determining differential expression in digital gene expression data using a model based on negative binomial distribution. The resulting P-values were adjusted using the Benjamini and Hochberg’s approach for controlling the false discovery rate. Genes with an adjusted P-value <0.05 found by DESeq were assigned as differentially expressed.

All data mentioned above were obtained from at least three replicates.

## Acknowledgements

This work was supported by National Key Research and Development Project of China No.2018YFA0901600 and National Natural Science Foundation of China No. 32070068.

