## Supplemental file for "An alternative malonyl-CoA producing pathway in nature"

***Supplementary S1. purification of BkdFGH complex***

For purification of the complex, $His_6$-tag was linked to the C-terminal of BkdF (E1α) to avoid affecting the assembly of E1 (β subunit C-terminal) and E2 core (subunit binding domain) [25], and the complex was expressed using the pSB1s plasmid (pSB1s-$His_6$-OADH) with the $P_{BAD}$ promoter. Our first attempts were unsuccessful because of the high expression level from the plasmid and incorrect subunit composition (for the FPLC fraction curve see Supplementary Fig. S1), resulting in erroneous assembly of the large protein complex, as previously described for other similar heterologous complex expression [26]. The expression cassette was then integrated to the *E. coli* chromosome *fadR* loci and the promoter was replaced by the constitutive medium-strength promoter $P_{tac}$ (strain His-OADH). N-terminal $His_6$-tagged E3 was purified separately using the pSB1s plasmid (pSB1s-$His_6$-LpdA1) considering that this component binds to the E2 core noncovalently and can dislocate from the complex [27].

SDS-PAGE analysis of the purified branched α-chain keto acid dehydrogenase revealed protein bands with molecular weights of 65 kDa, 50 kDa, 45 kDa, and 38 kDa, among which the molecular weight of BkdH is larger than the calculated value (49 kDa, see Supplementary Fig. S1). The bands were ascribed to BkdH, LpdA1, BkdF, and BkdG, respectively, using LC-MS/MS protein identification (Fig. 1g). The E3 component, to which the E2 component binds noncovalently [27], was dislocated from the complex upon further FPLC purification (Fig. 1g, Supplementary Fig. S1), revealing the weak interactions between the E2 and E3 components. Therefore, $His_6$-tagged E3 was added to OADH-$His_6$-tag at a weight ratio of 1:2 to obtain the entire complex just prior to assaying.

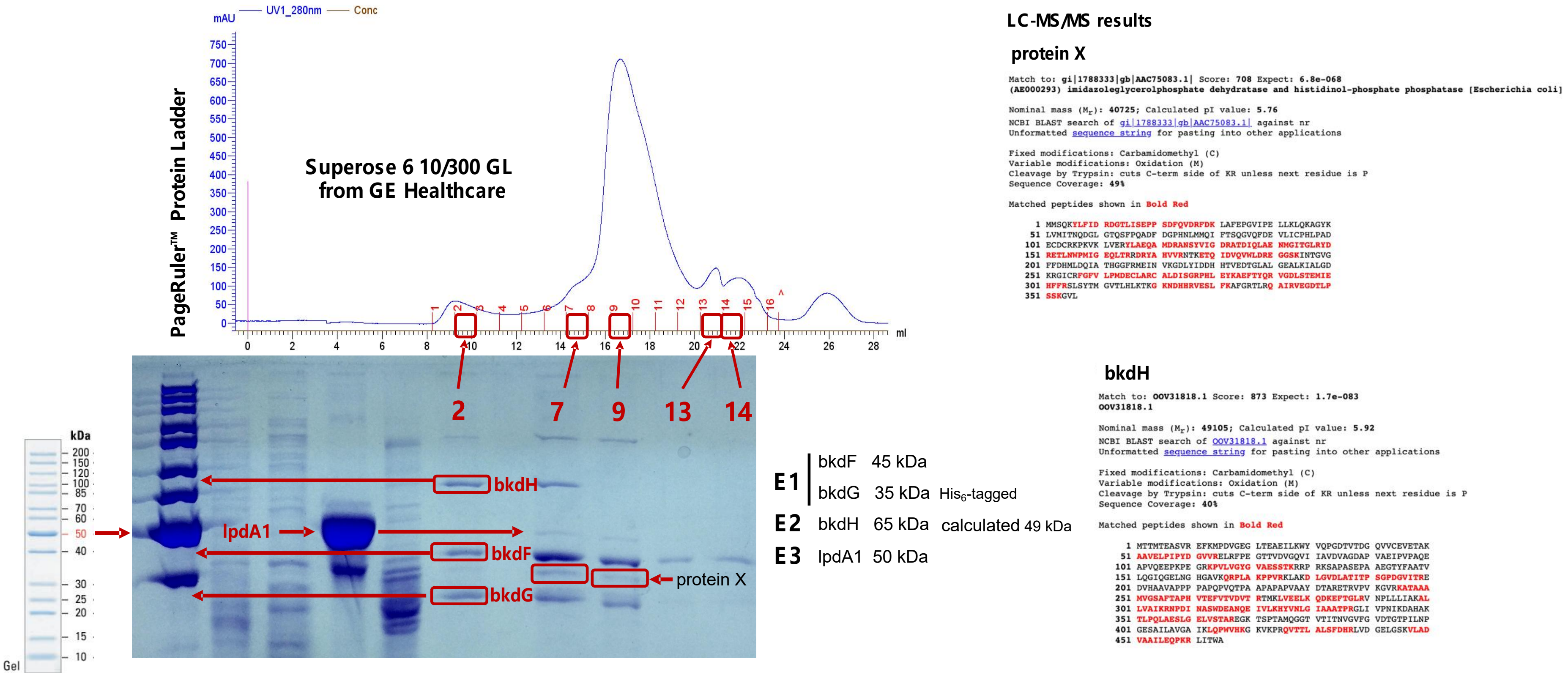


**Fig. S1. Complex purification by FPLC and protein verification by LC-MS/MS.**

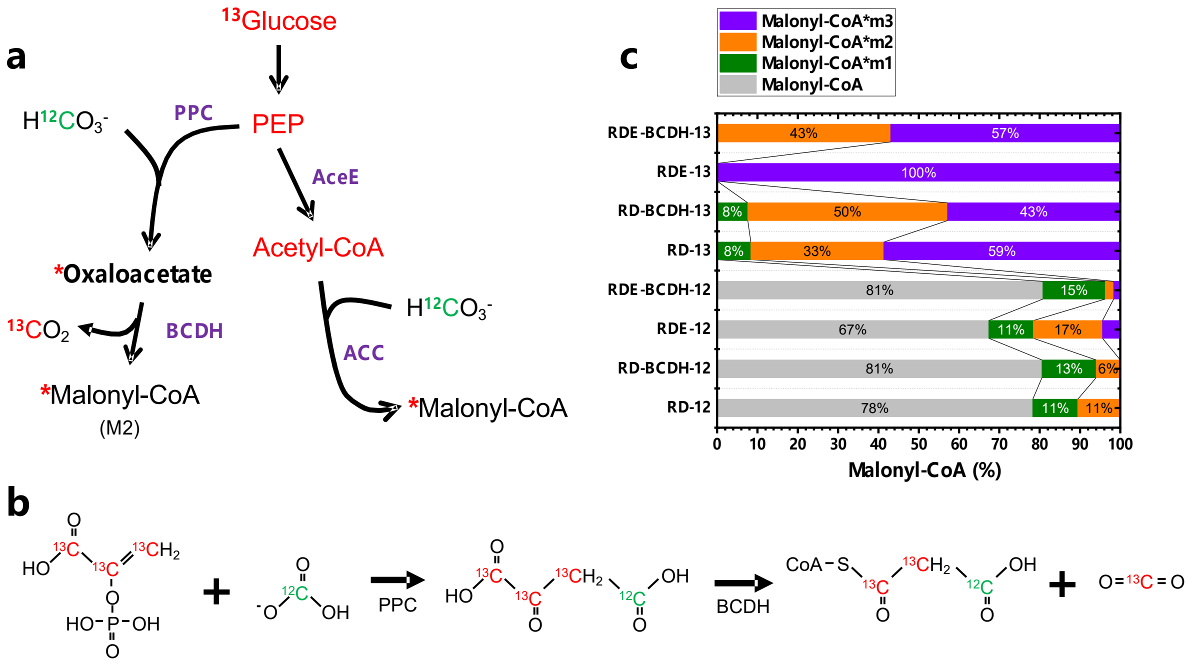


**Fig. S2 $^{13}$C- trace of malonyl-CoA in different biosynthesis pathway. a,** Two biosynthesis pathways of strain KDH. **b,** $^{13}$C-labeling positions in malonyl-CoA biosynthesis pathway via oxaloacetate. **c,** Proportion of malonyl-CoA with different numbers of $^{13}$C labelled. m1, m2 and m3 represent 1, 2 and 3 C atoms-labelled, respectively. RD: Δ*fadR*-Δ*panD*, RD-KDH: Δ*fadR*:: $P_{119}$-*bkdFG-bkdH-rbs-lpdA*-ΔpanD, RDE: Δ*fadR*-Δ*panD*-Δ*aceE*. RDE-KDH: Δ*fadR*::$P_{119}$-*bkdFG-bkdH-rbs-lpdA*-Δ*panD*-Δ*aceE*. 12 and 13 represent the strains were treated with [U-$^{12}C_6$] Glucose and [U-$^{13}C_6$] Glucose, respectively.

**Table S1. Strains, plasmids and promoter sequences used in this study.**

| Strain | Genotype | Source |
|---|---|---|
| *E. coli* BW25113/F′ | *rrnBT*14 Δ*lacZWJ*16 *hsdR*514 Δ*araBADAH*33 Δ*rhaBADLD*78 [F *proAB lacIqZ*ΔM15 *Tn*10 (*Tetr*)] | Coli Genetic Stock Center |
| Control | *E. coli* BW25113/F′, Δ*fadR* | This study |
| KDH | *E. coli* BW25113/F′, Δ*fadR*::$P_{119}$-*bkdFG-bkdH-rbs-lpdA1* | This study |
| KDH-Δ*ilvE* | Strain KDH, Δ*ilvE* | This study |
| KDH-ppc | Strain KDH, Δ*ompT*::$P_{tac}$-Cg-ppc | This study |
| ΔpanD | *E. coli* BW25113/F′, Δ*fadR,* Δ*panD,* carrying pS95s-GFP | This study |
| ΔpanD-KDH | Strain BCDH, Δ*panD,* carrying pS95s-GFP | This study |
| ΔpanD-mcrC | *E. coli* BW25113/F′, Δ*fadR,* Δ*panD,* carrying pS95s-MCR-C | This study |
| ΔpanD-mcrC-KDH | *E. coli* BW25113/F′, Δ*fadR*::$P_{119}$-*bkdFG-bkdH-rbs-lpdA1,* Δ*panD,* carrying pS95s-MCR-C | This study |
| ΔE1α | *E. coli* BW25113/F′, Δ*fadR*::$P_{119}$-*bkdG-bkdH-rbs-lpdA1,* Δ*panD,* carrying pS95s-MCR-C | This study |
| ΔE1β | *E. coli* BW25113/F′, Δ*fadR*::$P_{119}$-*bkdF-bkdH-rbs-lpdA1,* Δ*panD,* carrying pS95s-MCR-C | This study |
| ΔE2 | *E. coli* BW25113/F′, Δ*fadR*::$P_{119}$-*bkdFG-rbs-lpdA1,* Δ*panD,* carrying pS95s-MCR-C | This study |
| Δlpd | *E. coli* BW25113/F′, Δ*fadR*::$P_{119}$-*bkdG-bkdH,* Δ*panD,* carrying pS95s-MCR-C | This study |
| His-OADH | *E. coli* BW25113/F′, Δ*fadR*::$P_{119}$-$His_6$-*bkdF-bkdG-bkdH-rbs-lpdA1* | This study |
| RD | *ΔfadR-ΔpanD* | This study |
| RD-KDH | Strain KDH, Δ*panD-* | This study |
| RDE | Strain RD, Δ*aceE* | This study |
| RDE-KDH | Strain RD-KDH, Δ*aceE* | This study |
| Acc | *E. coli* BW25113/F′, Δ*fadR*::$P_{119}$-*accBC-rbs-accD-rbs-accA* | This study |
| Control-3HP | Strain ΔfadR carrying pSB1s-MCR-CN | This study |
| Acc-3HP | Strain Acc carrying pSB1s-MCR-CN | This study |
| OADH-3HP | Strain KDH carrying pSB1s-MCR-CN | This study |
| Control-TAL | Strain ΔfadR carrying pXB1k-PS1 | This study |
| Acc-TAL | Strain Acc carrying pXB1k-PS1 | This study |
| OADH-TAL | Strain KDH carrying pXB1k-PS1 | This study |

| | | |
|---|---|---|
| Control-Phl | Strain ΔfadR carrying pXB1k-PhlD | This study |
| OCDH- Phl | Strain KDH carrying pXB1k-PhlD | This study |
| Control-Aloesone | Strain ΔfadR carrying pLB1s-ALS | This study |
| OADH-Aloesone | Strain KDH carrying pLB1s-ALS | This study |
| Control-Flaviolin | Strain ΔfadR carrying pLB1s-RppA | This study |
| OADH-Flaviolin | Strain KDH carrying pLB1s-RppA | This study |
| Control-RK | Strain ΔfadR carrying pYB1a-BBCL | This study |
| OADH-RK | Strain KDH carrying pYB1a-BBCL | This study |
| Control-RVT | Strain ΔfadR carrying pAB1a-SCL | This study |
| OADH-RVT | Strain KDH carrying pAB1a-SCL and pYB1s-OADH | This study |
| RD-ppu-BCDH | *E. coli* BW25113/F$'$, Δ*fadR*::$P_{119}$-*bkdA1-bkdA2-bkdB-lpdV, ΔpanD* | This study |
| ΔpanD-mcrC-ppu-BCDH | *E. coli* BW25113/F$'$, Δ*fadR*::$P_{119}$-*bkdA1-bkdA2-bkdB-lpdV, ΔpanD,* carrying pS95s-MCR-C | This study |

| **Plasmid** | **Description** | **Source** |
|---|---|---|
| pSL91k | 119 promoter, pSC101 ori, Kan$^r$, selective marker flanked by lox71 and lox66 | Our laboratory |
| pSB1s | araBAD promoter, pSC101 ori, Str$^r$ | Our laboratory |
| pXB1k | araBAD promoter, p15A ori, Kan$^r$ | Our laboratory |
| pLB1s | araBAD promoter, R6k ori, Str$^r$ | Our laboratory |
| pYB1a | araBAD promoter, high-copy-number p15A ori variant, Amp$^r$ | Our laboratory |
| pAB1a | araBAD promoter, ColA ori, Amp$^r$ | Our laboratory |
| pYB1s | araBAD promoter, high-copy-number p15A ori variant, Str$^r$ | Our laboratory |
| pSL91k-BCDH | pSL91k containing *bkdFG-bkdH-rbs-lpdA1* | This study |
| pSL91k-ACC | pSL91k containing *accB-accC-rbs-accD-rbs-accA* | This study |
| pCP20 | Temperature sensitive vector carrying FLP recombinase, Amp$^r$ | [1] |
| pKD46 | Temperature sensitive vector carrying Red recombinase, Amp$^r$ | [1] |
| pSB1s*-Cre | Temperature sensitive vector carrying Cre recombinase gene, araBAD promoter, Str$^r$ | Our laboratory |

| | | |
|---|---|---|
| pS95s-MCR-C | 119 promoter, pSC101 ori, $Str^r$, expressing malonyl-CoA reductase | This study |
| pS95s-GFP | 119 promoter, pSC101 ori, $Str^r$, expressing green fluorescent protein | Our laboratory |
| pSB1s-$His_6$-BCDH | pSB1s containing $His_6$-*bkdF-bkdG-rbs-bkdH-lpdA1* | This study |
| pSB1s-$His_6$-LpdA1 | pSB1s containing $His_6$-*lpdA1* | This study |
| pSB1s-MCR-CN | pSB1s containing $mcr_{550\text{-}1219}$ N940V K1106W S1114R-rbs-$mcr_{1\text{-}549}$ | [2] |
| pXB1k-PS1 | pXB1k containing g2ps1 | This study |
| pXB1k-PhlD | pXB1k containing phlD | This study |
| pLB1s-ALS | pLB1s containing Rpa-ALS | This study |
| pLB1s-RppA | pLB1s containing RppA | This study |
| pYB1a-BBCL | pYB1a containinging BAR-BAS-4CL | This study |
| pAB1a-SCL | pAB1a containinging SLS-4CL | This study |
| pYB1s-OADH | pYB1s containinging *bkdFG-bkdH-rbs-lpdA1* | This study |

| **Promoter** | **Sequence** | **Source** |
|---|---|---|
| 119 | TTGACAGCTAGCTCAGTCCTAGGTATAATGCTAGCA | igem.org |

*Acetyl CoA carboxylase and actinorhodin biosynthesis genes annotation*

SCO0546: pyruvate carboxylase; SCO2445: acetyl CoA carboxylase subunits alpha/beta; SCO2776: acetyl/propionyl CoA carboxylase subunit beta; SCO2777: acetyl/propionyl CoA carboxylase subunit alpha; SCO4380: acetyl/propionyl CoA carboxylase subunit beta; SCO4381: acetyl/propionyl CoA carboxylase subunit alpha; SCO4921: acyl-CoA carboxylase complex alpha subunit; SCO4926: propionyl-CoA carboxylase complex B subunit; SCO5535: acetyl CoA carboxylase beta subunit; SCO5536: acetyl CoA carboxylase ε subunit; SCO6271: acyl-CoA carboxylase complex alpha subunit; SCO6284: acyl-CoA carboxylase complex alpha subunit. SCO5082: transcriptional regulator; SCO5083: actinorhodin transporter; SCO5085: actinorhodin operon activator protein; SCO5086: ketoacyl reductase; SCO5087: actinorhodin polyketide beta-ketoacyl synthase subunit alpha; SCO5088: actinorhodin polyketide beta-ketoacyl synthase subunit beta; SCO5089: actinorhodin polyketide synthase; SCO5090: actinorhodin polyketide synthase bifunctional cyclase/dehydratase; SCO5091: cyclase; SCO5092: actinorhodin polyketide dimerase.

*Isotopic labeling metabolic flux analysis*

$^{13}C$-trace experiments were conducted to verify the reaction as supplementary. Oxaloacetate was mainly replenished from PEP carboxylation in *Escherichia coli* [3]. Assuming malonyl-CoA was produced by oxaloacetate decarboxylation, [1,2-$^{13}C$] malonyl-CoA (M2) will be generated when the cell was treated with [U-$^{13}C_6$] Glucose and excess $NaH^{12}CO_3$ (Fig 5AB). As expected, malonyl-CoA with two acyl-C atoms (M2) labeled was increased from 33% (RD-13) to 50% (RD-BCDH-13) when the BCDH was overexpressed (Fig 5C). *aceE* was knockout (RDE) to lower intracellular acetyl-CoA level and block acetyl-CoA carboxylase pathway, the primary malonyl-CoA source of *Escherichia coli*. Only malonyl-CoA with three acyl-C atoms labeled (M3) was detected, while the proportion of M2 was increased significantly from undetectable to 43% (Fig 5C) when the complex was overexpressed (RDE-BCDH-13). Increased proportion of M2 by BCDH illustrated that malonyl-CoA was produced by oxaloacetate decarboxylation.

1. Datsenko, K.A. and B.L. Wanner, *One-step inactivation of chromosomal genes in Escherichia coli K-12 using PCR products.* Proceedings of the National Academy of Sciences, 2000. **97**(12): p. 6640-6645.
2. Liu, B., et al., *Efficient production of 3-hydroxypropionate from fatty acids feedstock in Escherichia coli.* Metabolic engineering, 2019. **51**: p. 121-130.
3. Kameshita, I., K. IZUI, and H. KATSUKI, *Phosphoenolpyruvate carboxylase of Escherichia coli.* The Journal of biochemistry, 1979. **86**(1): p. 1-10.
